# Poxviruses induce a nuclear environment in the cytoplasm to promote viral replication

**DOI:** 10.1101/2024.11.21.624458

**Authors:** Pia Veratti, Emanuel Wyler, Clarissa Read, Myriam Scherer, Jiang Tan, Paul Walther, Stephanie Wider, Daniel Weidl, Florian Full

**Author notes:** Correspondence to: Florian Full, PhD Institute of Virology, University Medical Center Freiburg, 79104, Freiburg, Germany.

## Abstract

The family Orthopoxviridae comprises several members that can cause disease in humans, including Small-, Cow-, Camel-, and Monkeypoxvirus. In 2022 a global outbreak of the zoonotic monkeypoxvirus started with human- to- human transmission and more than 100 000 global cases in all continents. This outbreak emphasizes the need for more basic research to get a better understanding of poxvirus biology.

We characterized the cellular response to poxvirus infection and performed bulk RNA-sequencing of primary human fibroblasts infected with Modified-Vaccinia Ankara Virus (MVA), Vaccinia Virus, Cowpoxvirus, Camelpoxvirus and 4 different Monkeypoxvirus isolates (clade 1-2b) at various time points to compare the transcriptome of infected cells. We could demonstrate that MVA infection leads to a potent activation of innate immune genes that is absent in other samples. Camelpoxvirus infection results in a strong activation of NF-kB signaling 2h post infection whereas all other viruses inhibit NF-kB activation. In addition, we observed transport of cellular proteins like histone H1, lamin A/C, emerin and topoisomerase II A/B from the nucleus into cytoplasmic poxvirus factories. This correlates with drastic structural changes inside the nucleus in electron microscopy, namely formation of nuclear filaments, indicating intensive remodeling of the nuclear architecture. Accordingly, inhibition of topoisomerase using the clinically approved topoisomerase inhibitor epirubicin reduced viral replication of several Mpox strains. We observe a reduction of viral replication by epirubicin in cells of human and mouse species, indicating that the recruitment of TOP2B into viral factories is crucial for MPXV replication.

## Introduction

The family of orthopoxviridae consists of a number of viruses that can cause disease in humans. Smallpoxvirus was among the deadliest diseases in human history until the WHO started a global vaccination program that ended with the worldwide eradication of smallpox in 1980. Since then, the zoonotic monkeypoxvirus (MPXV) which causes Mpox disease emerged as the most important orthopoxvirus of public health importance. For the past decades, there has been a continuous increase in MPXV cases, mostly restricted to Africa and caused by zoonotic spillover events ^1^. In 2022 a global outbreak of the zoonotic MPXV with a clade 2b virus started with human- to- human transmission and more than 100,000 global cases worldwide ^1,2^. In 2024 a novel clade 1b virus emerged in central Africa causing thousands of cases with hundreds of deaths and first cases were reported outside of Africa. As a consequence, the WHO declared a global health emergency in August 2024 ^3^.

Until the global outbreak in 2022, mpox was believed to have limited spread via human-to-human contact ^1^. The natural reservoirs of MPXV are probably small African rodents, but evidence remains sparse. Recent genomic data from the Democratic Republic of Congo (DRC) shows that alongside the current clade Ib epidemic, in recent years there have been multiple independent zoonotic transmissions of MPXVs across the country ^4^. It has been hypothesized that the current clade Ib outbreak partially results from such repeated spill-over events, which increase the likelihood of an emerging variant with better onward transmissibility among humans. Other zoonotic orthopoxviruses that are known to cause disease in humans are Camelpox virus and Cowpox virus. After the smallpox eradication, poxvirus research was neglected compared to research on other virus families and in the case of MPXV also hampered by its classification as biological select agent in the US and concerns about dual use. Existing data on the molecular biology of poxviruses is based on established techniques and most studies in the past years focused on the prototypic Vaccinia Virus (VAVC) ^5^. In particular, comparative “omics” approaches with different orthopoxviruses are missing. This and the current outbreak emphasize the need for more basic research to get a better understanding of poxvirus biology.

In contrast to other DNA viruses that replicate in the nucleus, poxvirus replication takes place in viral replication compartments (viral factories) in the cytoplasm ^5^. Upon infection of the cytoplasm, orthopoxvirus particles contain a transcription machinery that synthesizes viral mRNAs. Viral mRNA synthesis is detected within 20 minutes post infection ^6^. VACV transcription can be subdivided into three different types of transcripts, early, intermediate and late transcripts ^6^. Early transcription takes place between 20min and 2.5h post infection (p.i.). Early viral mRNAs are capped and polyadenylated like eukaryotic mRNAs, and the switch from early to intermediate transcription is mediated by viral decapping enzymes D9R and D10R that decap and thereby induce degradation of viral early mRNAs ^7,8^. Polyadenylation of viral mRNAs is mediated by viral proteins VP55 and VP39 ^9,10^. Intermediate transcription starts after viral DNA replication is finished - about 1.5h p.i. - and continues until late transcription starts at about 3h p.i.. Once late transcription is initiated, it continues until the end of the viral replication cycle. In addition, poxviruses induce a robust host shut-off on the host transcription and translation. The transcriptome of cells infected with a single poxvirus had been investigated by several labs, mainly for the prototypic VACV^11–18^. The goal of this study was a comparative analysis of the viral and host transcriptome in cells infected with eight different orthopoxviruses that can infect humans, in order to shed new light into the complex interactions between orthopoxviruses and the human host.

## Methods

### Cell culture and viruses

HFF1 cells (ATCC), BHK-21 cells (ATCC), TRF (Tert-immortalized rhesus fibroblasts, kindly provided by Dr. Philipp Kolb, Freiburg) and Vero E6 cells (ATCC) were cultured in Dulbecco’s modified Eagle’s medium (DMEM, Thermo Fisher Scientific) supplemented with 10% heat-inactivated fetal bovine serum (FBS, Anprotec), and 1% penicillin-streptomycin (Sigma-Aldrich). All cells were kept under standard culture conditions and monthly checked for mycoplasma contamination (MycoAlert ® Mycoplasma detection Kit, Lonza).

Infection experiments were performed with Vaccinia Virus–Western Reserve (VACV-WR), Modified Vaccinia Ankara (MVA), Monkeypoxvirus-Gabon (MPXV-GA, Gabon-1987, Clade IIa), Monkeypoxvirus-Ivory Coast (MPXV-IC, Ivory Coast-2012, Clade I), Monkeypoxvirus-Freiburg (MPXV-FR, Freiburg-2022, Clade IIb), Monkeypoxvirus-Robert Koch Institute (MPXV-RKI, RKI-2022, Clade IIb), Cowpoxvirus (CPXV, Hum/Gri 07/01) and Camelpoxvirus (CMLV, CP-1).

### Virus infection

For RNA-Seq and RT-qPCR experiments HFF1 cells were seeded in 6 well plates the day before infection. Before infection, an aliquot of the cell supernatant was removed and kept at 37°C. Cells were infected at an MOI=1 and after 2h incubation at 37°C the virus-containing medium was replaced by the cell supernatant kept previously.

For immunofluorescence experiments HFF-TERT or TRF cells were seeded on cover glasses in 24 well plates the day before infection. Cells were infected at an MOI=0,5.

For inhibitor treatment experiments HFF-TERT or TRF cells were seeded in 24 well plates the day before infection. Cells were treated with either the inhibitors or DMSO and at the same time infected at an MOI=0,5. After 2h incubation at 37°C, the virus-containing supernatant was replaced by fresh medium containing the inhibitors or DMSO (Carl Roth).

### Next generation RNA-Sequencing

HFF1 cells (ATCC) were infected with indicated orthopoxviruses at MOI of 1. At 2, 6 and 12hpi, cells were lysed in Trizol (Life Technologies by Thermo Fisher Scientific), and total RNA was isolated using the Direct-zol RNA Miniprep kit (Zymo Research), according to the manufacturer’s instructions. Poly-adenylated RNAs were enriched with the NEBNext Poly(A) mRNA Magnetic Isolation Module (NEB) and sequencing libraries were prepared using the NEBNext Ultra II Directional RNA Library Prep Kit for Illumina (NEB) according to the manufacturer’s instructions, with 9 cycles PCR amplification. Pooled libraries were sequenced on a Novaseq 6000 device with 2x109 cycles.

### Immunofluorescence

Supernatant of infected cells was removed and after washing with PBS, cells were fixed for 5 min at RT with 4% PFA and washed again with PBS. Blocking and permeabilization was performed in PBS containing 5% FCS and 0,1% Triton X-100 (Sigma-Aldrich) for 1h at RT. Primary antibodies (Table 1) diluted 1:500 in blocking and permeabilization buffer were added to the cells and incubated 1,5h RT or overnight at 4°C. After washing with PBS, cells were incubated 1h RT in permeabilization and blocking buffer containing fluorescent secondary antibodies (1:500) and Hoechst 33342 (Sigma-Aldrich) (1:1000). After washing with PBS, samples were mounted in Mowiol mounting medium. Imaging was performed with a Zeiss LSM880 Airyscan and Zeiss ZEN 3.0 (Blue edition) software was used for image editing.

**Table 1.**

| Primary Antibodies |  |  |  |
| --- | --- | --- | --- |
| Anti- Emerin (H-12) | mouse monoclonal | Santa Cruz<br>Biotechnology Inc | sc-25284 |
| Anti- Histone H1 (AE-4) | mouse monoclonal | Santa Cruz<br>Biotechnology Inc | sc-8030 |
| Anti- Histone H3 (1G1) | mouse monoclonal | Santa Cruz<br>Biotechnology Inc | sc-517576 |
| Anti- Lamin A/C (636) | mouse monoclonal | Santa Cruz<br>Biotechnology Inc | sc-7292 |
| Anti- TOPO II $\beta$ (H-8) | mouse monoclonal | Santa Cruz<br>Biotechnology Inc | sc-25330 |
| Anti- TOPO II $\alpha$ (G-6) | mouse monoclonal | Santa Cruz<br>Biotechnology Inc | sc-166934 |
| Anti- Vaccinia Virus | rabbit polyclonal | Invitrogen | PA1-7258 |

### Electron microscopy

Samples for electron microscopy were prepared as described in Read et al.^19^ with some modifications. In brief, HFF1 cells were cultured on carbon-coated sapphire discs (diameter: 3 mm, thickness: 160 µm, Engineering Offices M. Wohlwend GmbH, Switzerland) for 24 h and infected with MVA for 8, 16 and 24 h or left uninfected. Cells infected with MVA were cyro-immobilized from the native state without aldehyde fixation.

For cryoimmobilization, cells on sapphire discs were high-pressure frozen in a Wohlwend HPF compact 01 high-pressure freezer (Engineering Office M. Wohlwend GmbH, Switzerland). Samples were subsequently freeze-substituted in a solution containing 0.2 % (v/v) osmium tetroxide (ChemPur, Germany) and 0.1% uranyl acetate (w/v) (Merck KGaA, Germany) in acetone, supplemented with 5% (v/v) water to enhance membrane contrast (Walther and Ziegler, 2002). Over a period of 17 hours, the temperature was gradually increased from −90°C to 0°C, with a one-hour incubation step at 0°C and a further increase to room temperature within one hour. Samples were then rinsed three times with acetone, infiltrated incrementally with Epon 812 (30%, 60%, and 100% Epon in acetone), and finally polymerized at 60°C for at least 48 hours. The sapphire discs were removed, leaving the cells and the carbon coat on the surface of the Epon block.

For transmission electron microscopy (TEM), 70 nm sections were cut using an Ultracut UCT ultramicrotome (Leica, Germany) equipped with a diamond knife (Diatome Ltd, Switzerland). Sections were collected on freshly glow-discharged single slot copper grids which were coated with a carbon-reinforced Formvar film. The sections were examined with a JEM-1400 transmission electron microscope (Jeol, Japan) operating at 120 kV acceleration voltage. Images were acquired using a Veleta CCD camera (Olympus, Japan).

### qRT-PCR

RNA was extracted using the Direct-zol RNA Miniprep Plus kit from Zymo Research according to the manufacturer’s instructions. Reverse transcription and qRT-PCR were conducted in one step using 33ng RNA as template with the Luna Universal Probe One-Step RT-qPCR kit (New England Biolabs) following the manufacturer’s protocol in a Biorad C1000 Touch system. Primers/probes are listed in Table 2. Expression levels for each gene were obtained by normalizing values to *HPRT1* and fold induction was calculated using the comparative CT method (ΔΔCT method).

**Table 2.**
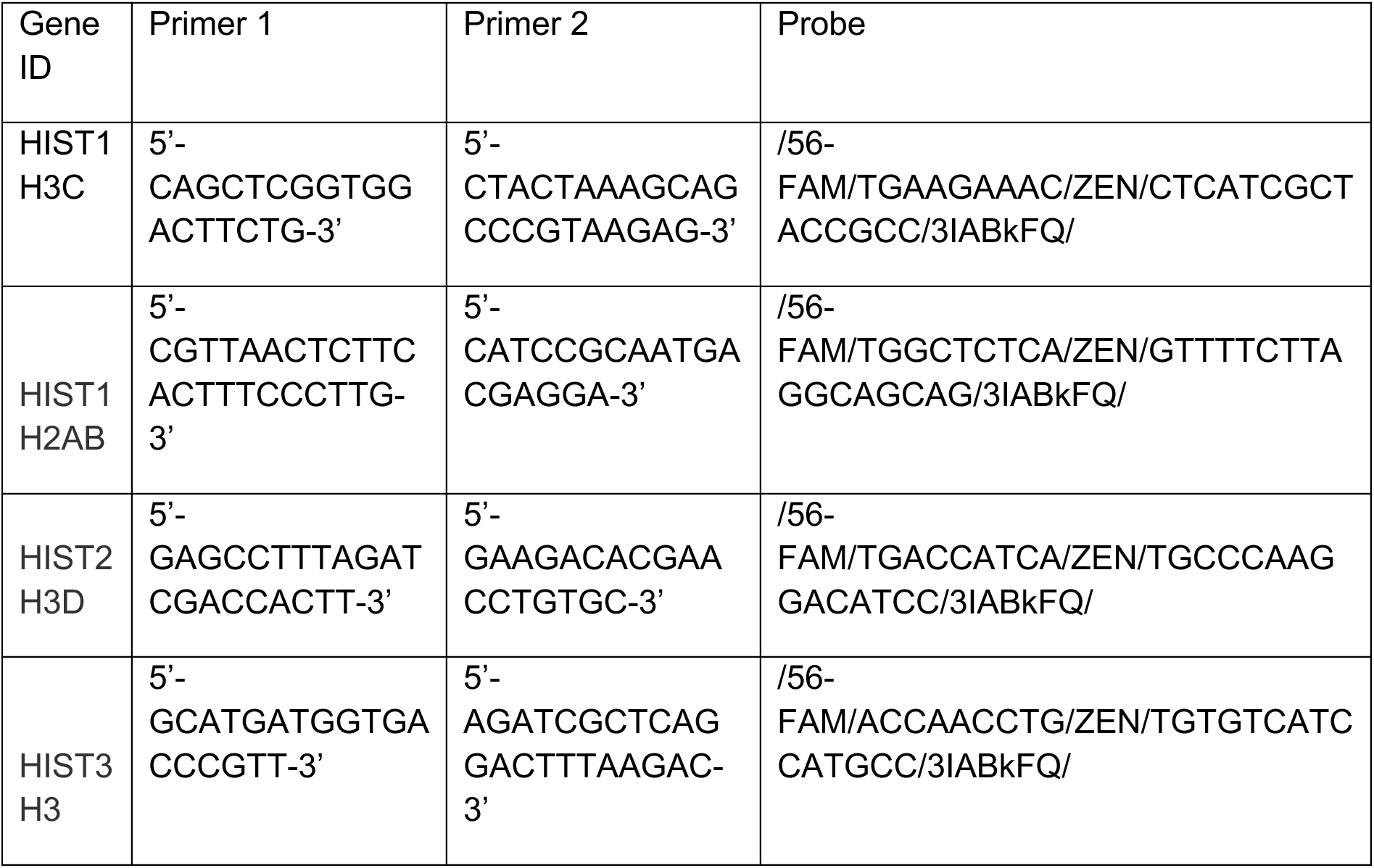
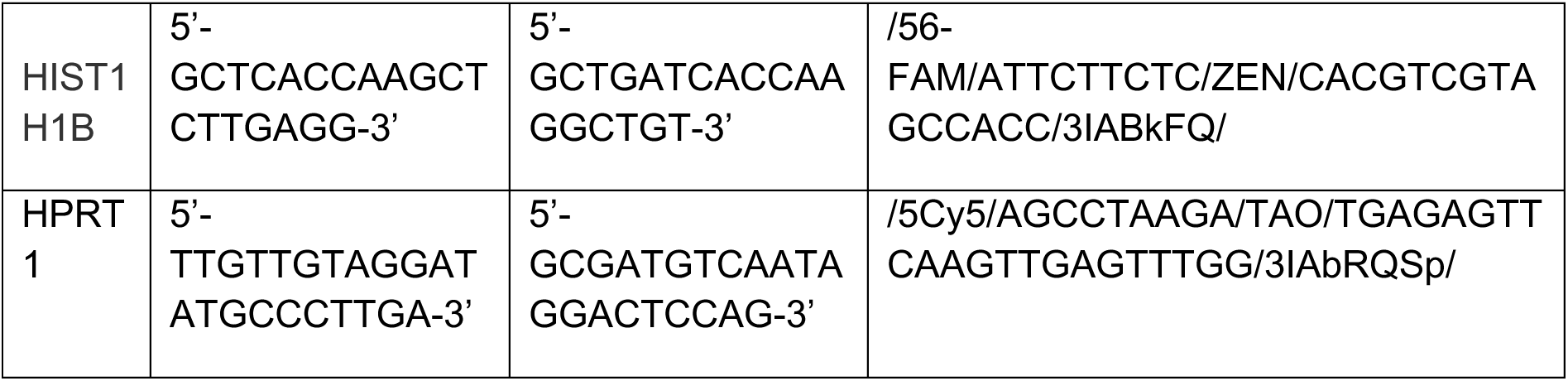

### Inhibitor treatment

Cells were treated in concomitance with the virus infection and 48 hours post infection by adding KPT-330 (Selinexor, SelleckChem) to a final concentration of 2µM or Epirubicin (Pfizer) to a final concentration of 0,2µM. The IC50 of both inhibitors was determined beforehand to choose a concentration that is well tolerated by cells.

### Plaque Assay

To determine the virus titer, Vero E6 cells were seeded in 12 well plates. Virus-containing supernatant was diluted at 10^−1 to 10^−4 in PBS and added to the cells. After incubating 1,5h at RT, supernatant was removed and cells coated with Oxoid agar solution (DMEM, 0,04% Oxoid Agar). The plaque number was determined 5 days post-infection by crystal violet staining.

## Results

In order to compare the transcriptome of cells with different poxviruses, we infected HFF1 cells with MVA, VV-WR, CPXV, CMLV, and four different monkeypoxvirus strains (1x clade 1a, 1x clade 2a and 2x clade 2b). Cells were harvested at 2, 6 and 12 hours post infection (h.p.i.) and performed bulk RNA sequencing (Figure 1A and B). We did not observe major differences in the expression kinetics of conserved orthopoxviruses genes for MVA, VV and MPXV. CPXV and CMLV seemed to replicate slower in human cells, as shown for example by delayed expression of early genes OPG147 (A44L) and OPG200 (B15R) (Fig. 1C, Fig. S1) compared to VACV and MPXV. For expression of cellular genes, there was an upregulation of JUN and FOS genes for all viruses except VACV and CPXV with slightly different kinetics (Fig. 1D), which indicates activation of the TNF pathway. The upregulation of JUN for example could be detected as early as 2h for CMLV, but started at 6h post infection for MPXV (Fig. 1D). We also observed an upregulation of EGR1 in MPXV infected cells, a transcription factor that has been shown to promote VACV replication ^20^. In addition, we observed a strong activation of the NF-kB signaling pathway in cells infected with CMLV which was completely absent for all other orthopoxviruses tested (Fig. 1E). Moreover, several heat shock proteins were upregulated upon infection with MPXV clade 2 viruses at the 12h time point, but not for all other viruses tested (Fig. 1F).

**Figure 1:**
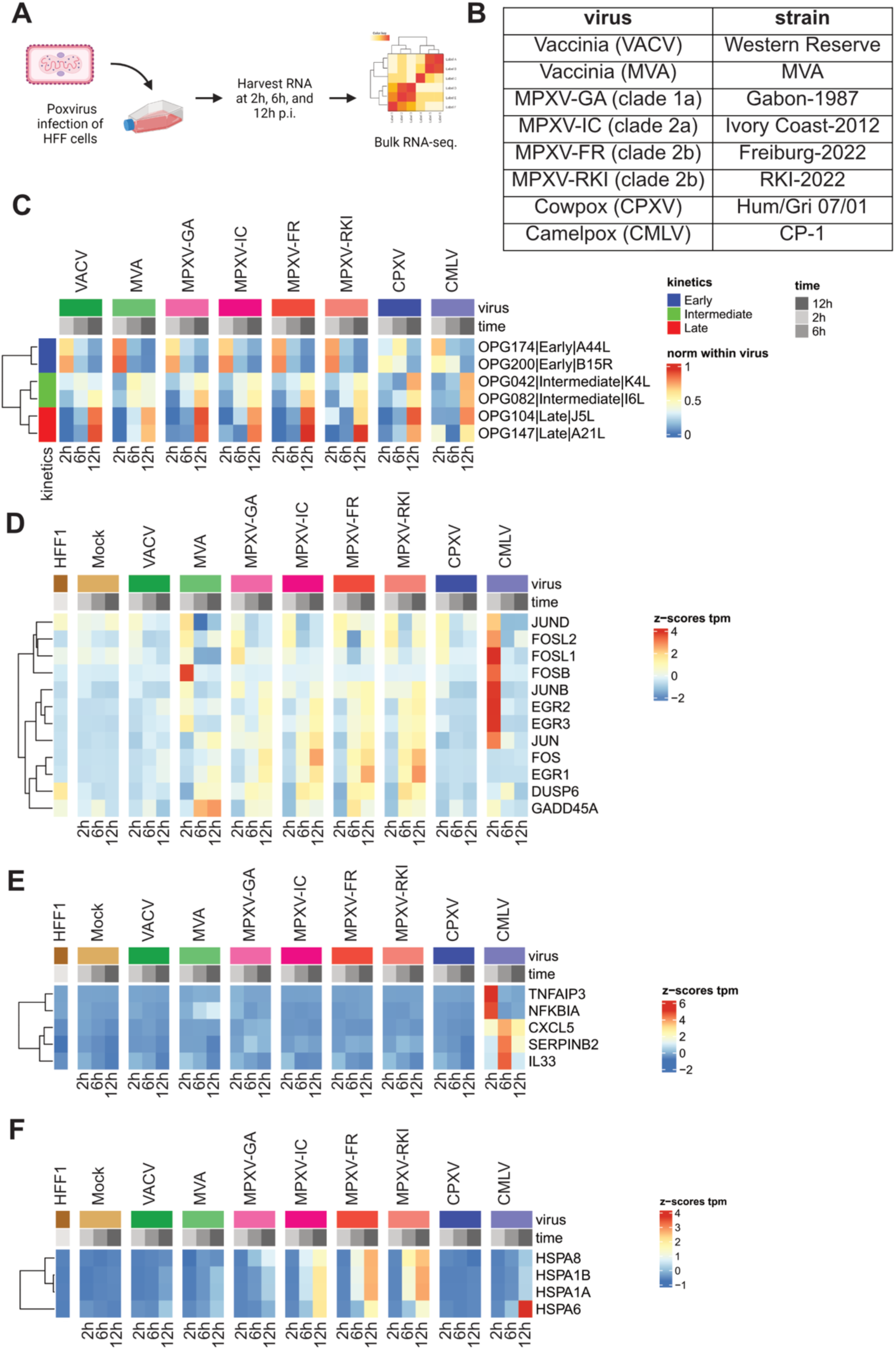
Comparative transcriptomics of orthopoxviruses. **A)** Schematic overview of the sequencing workflow. HFF1 cells (Human Foreskin Fibroblasts) were infected with MOI of 1 and harvested at indicated time points. **B)** Table of viruses used in the study. MPXV=Monkeypoxvirus; MVA=Modified Vaccina Ankara Virus **C)** Heatmap of expression of selected core orthopoxvirus genes with early, intermediate and late expression kinetics. **D)** Heatmap of expression of selected host genes of the TNF / MAPK pathways upon orthopoxvirus infection. **E)** Heatmap of expression of selected NF-kB pathway genes **F)** Heatmap of expression of different heat shock proteins.

Overall, the cellular response to MVA infection was different compared to the other orthopoxviruses in our panel, reflecting the prolonged cell culture adaptation in chicken cells. For example, there was no activation of the type I IFN pathway for all viruses tested except for MVA, which is a consequence of the potent immune evasion that orthopoxviruses are known for and that was lost in MVA (Fig. S2). We also observed a strong upregulation of cellular histone genes upon MVA infection which was less visible in MPXV infected cells and completely absent in VACV, CPXV and CMLV infected cells (Fig. 2A). To investigate whether this change in histone gene expression also results in changes in protein expression or localization, we performed immunofluorescence staining of infected HFF1 cells and stained for histone H1 (Fig. 2C). Whereas mock infected cells showed the expected nuclear expression of histone H1, we observed a localization of histone H1 to cytoplasmic viral factories upon infection with all orthopoxviruses in our panel, indicated by co-staining with a VACV polyclonal antibody. This was unexpected, because it was known from the literature that detection of non-polyadenylated histone mRNAs by poly-A based RNA-seq is rather a known consequence of poxviral poly-adenylation of transcripts by the viral poly(A) polymerase than true upregulation of mRNAs ^18,21^. We confirmed this by taqman qRT-PCR for several histone genes and could not detect any changes in the levels of histone mRNAs upon poxviral infection (Fig. 2C), indicating that the observed differences in histone mRNA levels in our RNA seq. were indeed due to a polyadenylation artifact of usually non-polyadenylated transcripts.

**Figure 2:**
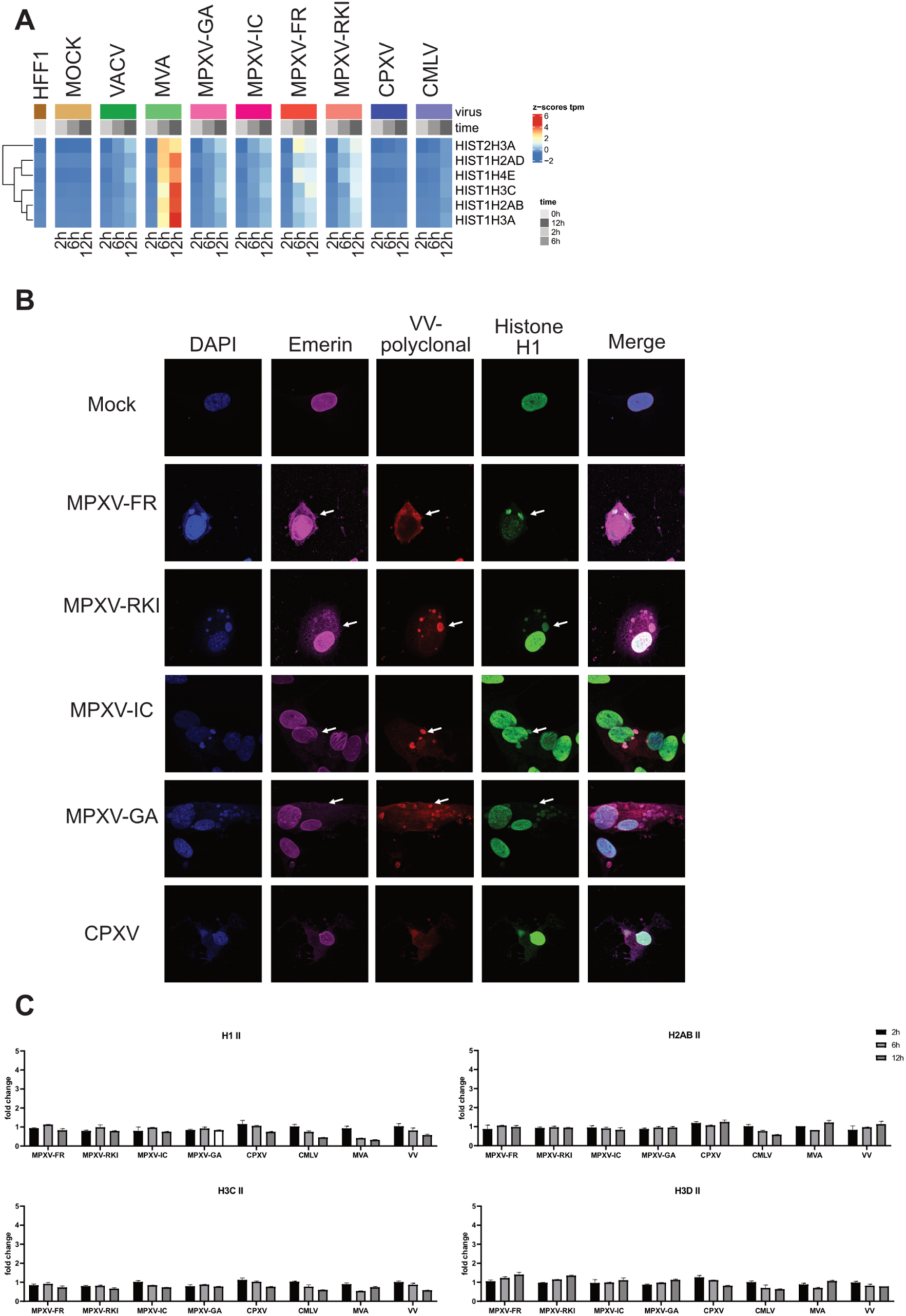
Histone 1 relocalization upon orthopoxvirus infection. **A)** Heatmap of expression of selected histone genes. **B)** Taqman qRT-PCR for histone variants H1II, H2ABII, H3CII and H3DII upon infection with our orthopoxvirus panel. The fold change was calculated relative to mock cells and the housekeeping gene HPRT1 (delta-delta Ct method). **C)** Immunofluorescence analysis of HFF cells infected with different MPXV strains and Cowpoxvirus. Cells were fixed and stained with antibodies for Emerin, Vaccinia Virus (polyclonal) and histone H1. Nuclei were counterstained with DAPI.

Next, we assessed whether the translocation of histone H1 into poxviral factories is a common mechanism for all histone proteins or even other nuclear proteins, we also stained infected cells for the nuclear lamina component emerin (Fig. 2B), histone H3 (H3, Fig. 3A), histone H2B (data not shown), Lamin A/C, topoisomerase 2A and topoisomerase 2B (Figure 4A). Whereas histone H3 and histone H2B showed no relocalization into viral factories upon infection, we observed colocalization of emerin, Lamin A/C, TOP2A and TOP2B with viral factories. This indicates a selective recruitment of cellular proteins into viral factories.

**Figure 3:** Lamin A/C relocalization upon orthopoxvirus infection. **A)** Immunofluorescence analysis of HFF cells infected with different MPXV strains, Cowpox (CPXV)- and Camelpoxvirus (CMLV). Cells were fixed and stained with antibodies for Lamin A/C, Vaccinia Virus (polyclonal) and histone H3. Nuclei were counterstained with DAPI.

**Figure 4:**
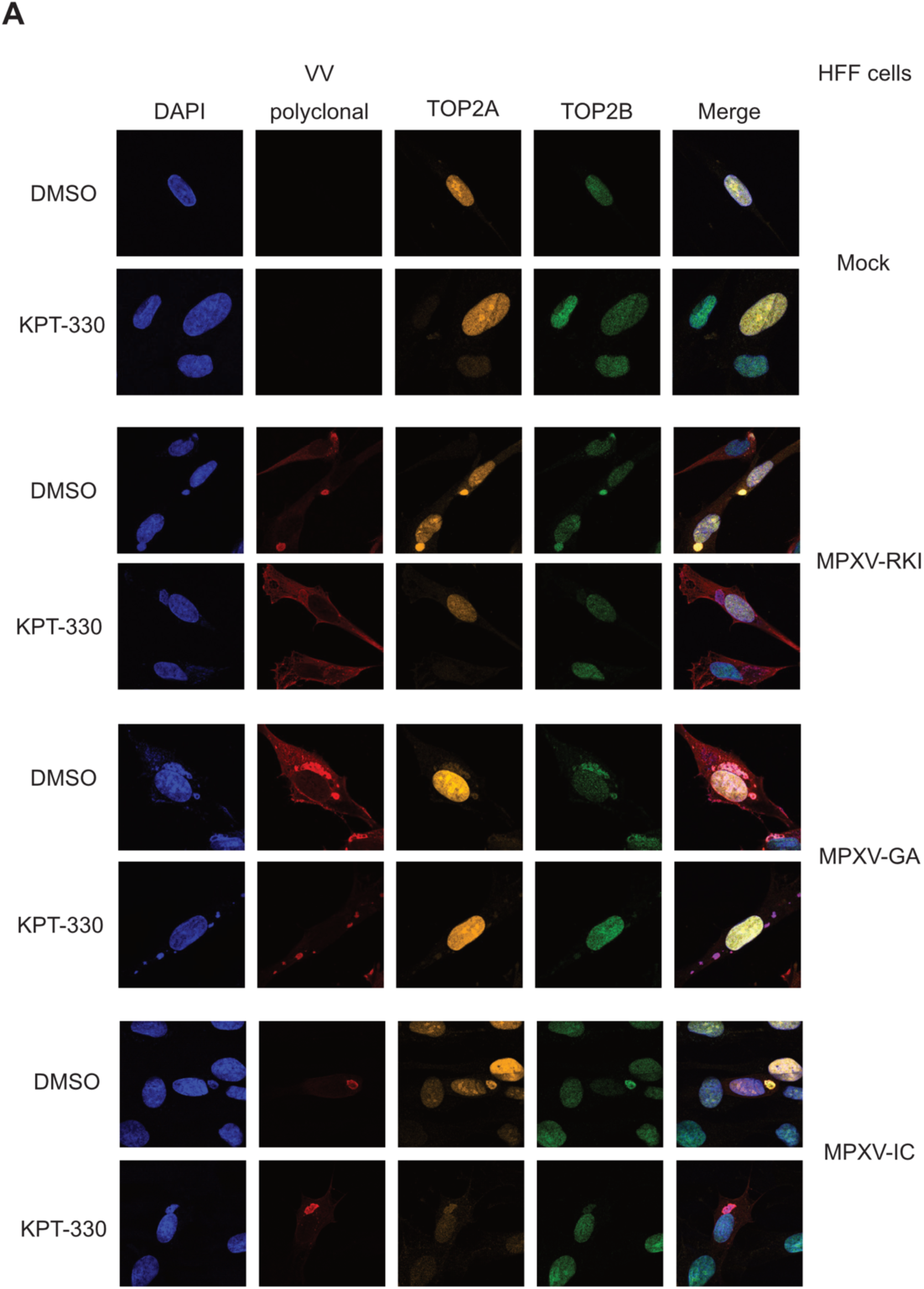
Cytoplasmic transport of Topoisomerase 2 subunits is dependent on the exportin-1 pathway. **A)** Immunofluorescence analysis of HFF cells infected with MPXV strain RKI (MPXV-RKI), MPXV strain GA (MPXV-GA) and MPXV strain Ivory Coast (MPXV-IC). Cells were either treated with DMSO (control) or the exportin-1 inhibitor KPT-330 (Selinexor). Cells were fixed and stained with antibodies for Vaccinia Virus (polyclonal), Topoisomerase 2A (TOP2A) and Topoisomerase 2B (TOP2B). Nuclei were counterstained with DAPI.

Selected nuclear proteins were detected in viral factories but it was not clear whether nuclear proteins are actively transported out of the nucleus into viral factories or whether newly synthesized proteins remain in the cytoplasm. The clinically approved exportin 1 inhibitor Selinexor was used to block export from the nucleus into the cytoplasm. Selinexor could efficiently block export of TOP2B into viral factories, which shows that nuclear export is responsible for the observed translocation in fibroblasts of human, rhesus and mouse origin (Fig. 4A, Fig. 5A, Fig. S5 and S6). In addition, electron microscopy of MVA infected cells revealed drastic changes in the nuclei of infected cells. We observed the formation of nuclear filaments of unknown origin in a fraction of infected cells that was completely absent in uninfected cells, indicating rearrangement of the nuclear architecture upon infection (Fig. 5B).

**Figure 5:**
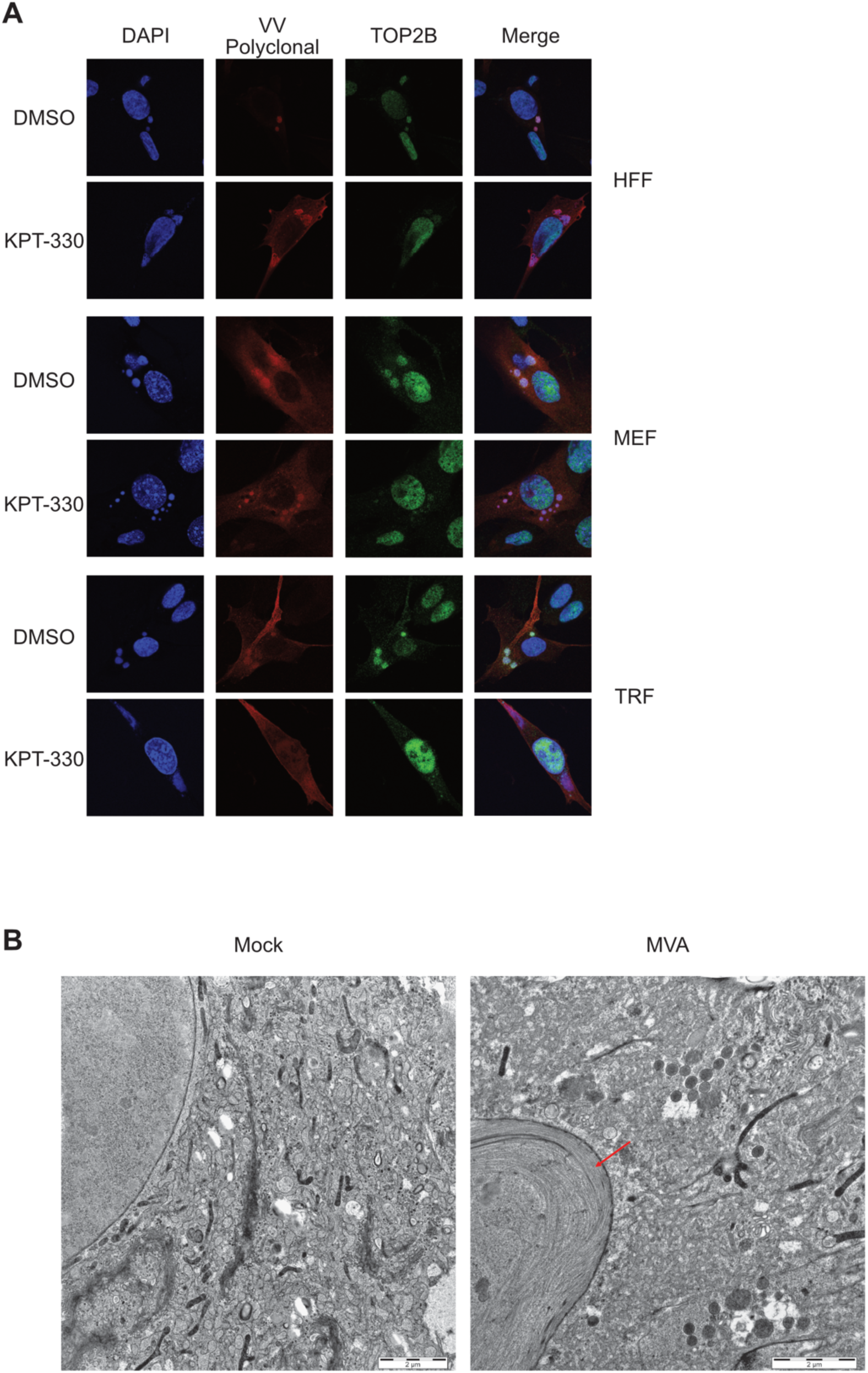
Cytoplasmic transport of Topoisomerase 2 subunits is dependent on the exportin-1 pathway. **A)** Immunofluorescence analysis of HFF cells, murine embryonal fibroblasts (MEF cells) and Tert-immortalized rhesus fibroblasts (TRF cells) infected with MPXV strain Freiburg (MPXV-FR). Cells were either treated with DMSO (control) or the exportin-1 inhibitor KPT-330 (Selinexor). Cells were fixed and stained with antibodies for Vaccinia Virus (polyclonal) and Topoisomerase 2B (TOP2B). Nuclei were counterstained with DAPI. **B)** Electron microscopy of HFF cells infected with Modified Vaccinia Ankara (MVA). 14hpi, unfixed cells were flash frozen in liquid nitrogen and subjected to EM analysis.

To investigate if the recruitment of TOP2 subunits is needed for viral replication, we treated infected cells with the clinically approved TOP2 inhibitor Epirubicin, which can either act as TOP2 poison or TOP2 inhibitor^22^. Using a nanomolar concentration that was shown to act as TOP2 poison, Epirubicin shows only minimal cytotoxicity in cells. The replication of all MPXV strains tested was impaired in human fibroblasts compared to DMSO treated control cells (Fig. 6A-D) by plaque assays of supernatants from infected cells. For MPXV infected murine fibroblasts we also observed a significant difference in infectious particles released from cells but to a lesser extent than in human HFF cells (Fig. 6A-D). This argues for the need of recruitment of TOP2 genes into viral factories and an important role of TOP2 in orthopoxvirus replication.

**Figure 6:**
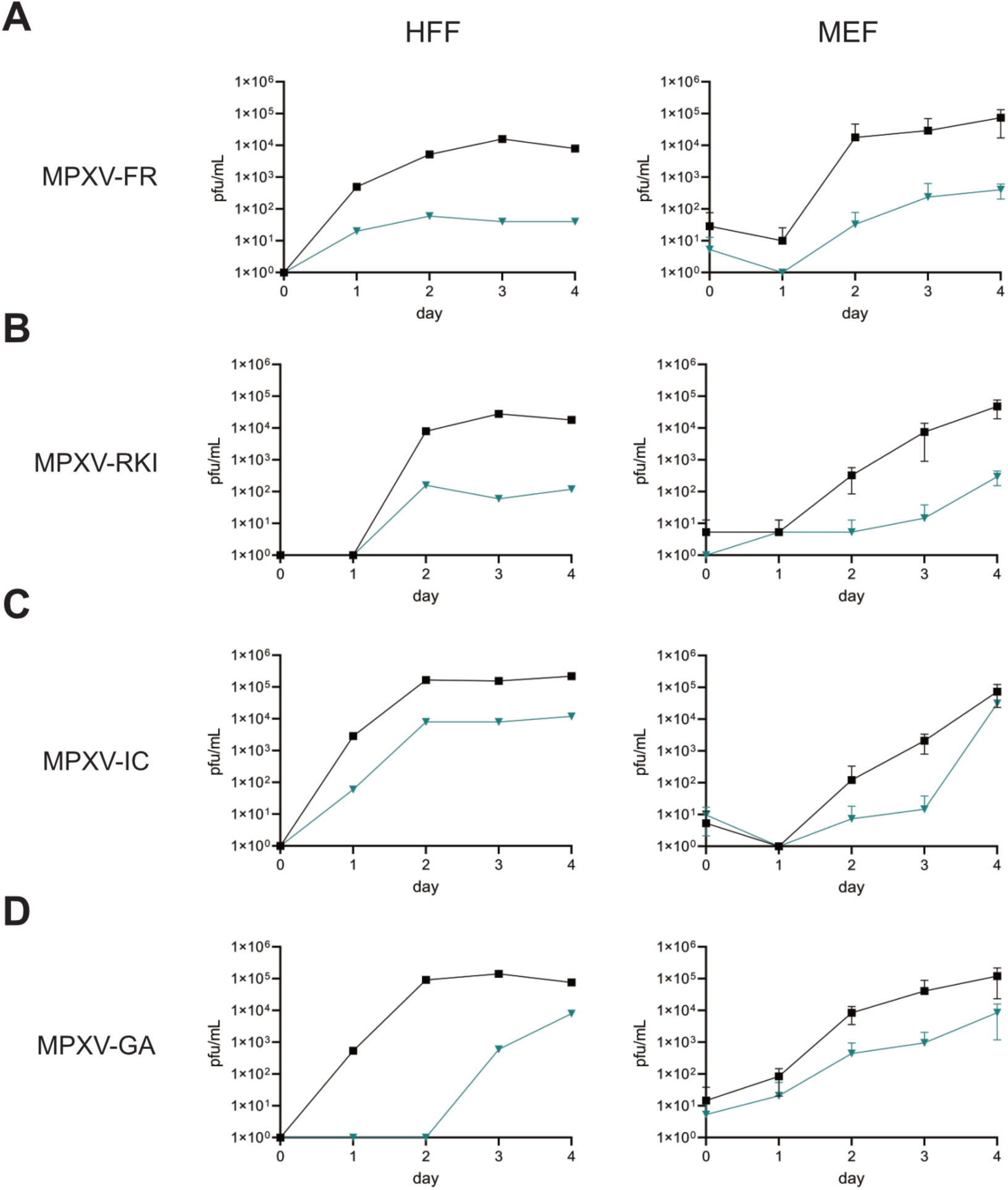
Topoisomerase inhibitor epirubicin impairs MPXV replication. Replication of different MPXV strains in HFF cells and MEF cells in the presence of DMSO (black line) and epirubicin (blue line). Cells were infected with MOI of 0.01 and release of infectious particles in collected supernatants (day 1-4) measured by plaque assay on Vero cells. **A)** MPXV-FR **B)** MPXV-RKI **C)** MPXV-IC **D)** MPXV-GA

## Discussion

Our study demonstrates that infection of several orthopoxviruses leads to the selective transport of nuclear proteins into poxviral factories, including TOP2A and TOP2B proteins. Poxvirus replication takes place in so called viral factories in the cytoplasm, in contrast to most DNA viruses that replicate in the nucleus. Cytoplasmic replication of DNA comes with a high price, because all enzymes that are needed for DNA replication have to be present in the cytoplasm, which is normally not the case. This is achieved by either being encoded by the virus itself, like the viral polymerase, or by transport of nuclear enzymes to the sites of replication. For example, it was reported before, that VACV needs proteins of the nuclear pore complex for efficient replication ^23^. In addition, it was also reported, that the nuclear proteins YY1, Sp1 and TATA-binding protein colocalize with viral replication compartments in the cytoplasm factories ^24^. However, the authors showed that export of YY1 could not be inhibited by Leptomycin B, another inhibitor of the exportin pathway. This points at a slightly different mechanism compared to our study, because transport of TOP2B into viral factories was inhibited by treatment with Selinexor, an inhibitor of the exportin 1 pathway. We also observed relocalization of histone H1 to viral factories in the presence of Selinexor (Fig. S3, Fig. S4), which can be explained by the small size of histone H1 and free diffusion through nuclear pores into the cytoplasm. In this regard, it would be of great interest to isolate poxvirus factories and nuclei from infected cells and perform comparative proteomics to analyze the change of protein localization upon infection in the presence and absence of various exportin inhibitors. But so far, we were not able to establish protocols to clearly separate nuclei from viral factories, as indicated by substantial amounts of infectious particles in the nuclear fractions. (data not shown). Of note, Matía et al. also observed transport of cellular proteins in between different organelles upon VACV infection ^17^.

The export of nuclear proteins correlates with the formation of nuclear filaments in MVA infected cells (Fig. 5B). Using electron microscopy, we observe the formation of nuclear filaments at later timepoints (24 hpi) in MVA infected cells. Unfortunately, the size and form of the filaments in electron microscopy did not provide any further insights into their nature. The formation of nuclear filaments was also observed *in vitro* and *in vivo* in cells infected with stomatitis papulosa virus (*<u>Parapoxvirus bovis 1</u>)* and Orf virus, two members of the genus Parapoxviruses that cause disease in cattle and sheep ^25^. In addition, chromatin conformation capture assays showed that VACV infection also results in drastic changes in the nuclear architecture ^20^. Taken together, this indicates that remodeling of the nucleus is a characteristic of poxvirus infection and we hypothesize that the export of nuclear proteins into viral factories either contributes to the observed nuclear alterations or is a consequence of the filament formation.

TOP2A and TOP2B are proteins that catalyze transient double strand breaks to relax supercoiled DNA and allow DNA replication. Poxviruses encode for their own Topoisomerase I gene (OPG 111) that catalyzes transient single-strand breaks^26^, but not for a Topoisomerase II gene. Topoisomerases are well known drug targets for both anti-bacterial and anti-tumor chemotherapy and the eukaryotic TOP2 inhibitors Etoposide, Doxorubicin and Epirubicin are widely used for combinational chemotherapy in the treatment of various tumors including liver-, bladder-, stomach-, breast- and lung-cancer ^27^. Although TOP2 inhibitors have considerable side effects in high dose treatment regimens for anti-tumor chemotherapy, they are also effective as so-called topoisomerase poisons in much lower concentrations ^22^. In vitro, epirubicin acts both as TOP2A poison and also as an inhibitor of TOP2B ^22^. We could show that epirubicin inhibits the replication of several MPXV strains of clades 1a, 2a and 2b in human and mouse fibroblasts in nanomolar concentrations (Fig. 6 A-C). We also observed transport of TOP2A and TOP2B into viral factories in rhesus fibroblasts, which indicates conservation of the mechanism between mammalian species (Fig. S5).

MPXV was initially isolated from crab-eating macaques in an animal facility in 1959 where it got its original name monkeypox virus from ^28^. This name is misleading since the natural reservoirs of MPXV are not monkeys but small rodents in Africa ^1^, however, the exact reservoir species has yet to be determined. MPXV has only been isolated from wild monkeys in two occasions, the first isolation was from a Thomas’s rope squirrel (Funisciurus anerythrus) in the Democratic Republic of the Congo ^29^. In addition, the monkeypoxvirus strain Ivory Coast (MPXV-IC) used in this study was isolated from a deceased infant sooty mangabee (Cercocebus atys) in Taï National Park in Ivory Coast in 2012 ^30^. MPXV-TNP is closely related to the strain isolated from a human in Liberia in 1970 and belongs to the West-African clade 2a of mpox strains ^31^. Patrono et al. reported MPXV infections in a chimpanzee population in Ivory Coast with several different MPXV strains over a period of 16 months ^31^. The chimpanzees had a variety of syndromes including respiratory illness without the characteristic skin rash. However, very little is known about MPXV disease in rhesus monkeys. Laboratory infections of related cynomolgus monkeys (Macaca fascicularis) with high dose (≥10,000 pfu) of MPXV-Zaire resulted in systemic infections and subsequent death ^32^. Our data point at a conserved mechanism in the need of TOP2 genes for viral replication since epirubicin was needed for MPXV replication in human and mouse fibroblasts. It would be very interesting to also test viral replication in cells of other mammalian species We hypothesize that the recruitment of nuclear proteins into viral factories is a critical step in the orthopoxvirus replication cycle that could be exploited for antiviral therapy, as exemplarily shown by the use of epirubicin.

## Data availability statement

Sequencing data (RNA-Seq.) were deposited in GEO with GEO accession number GSE234489.

## Acknowledgments

We are grateful to Andreas Nitsche (Berlin) and Toni Rieger (Hamburg) for providing virus stocks. We thank Aura M Bastidas-Quintero for her help in graphical design. This study was supported by grants from the BMBF Junior Research Group “Duxdrugs” (BMBF 01KI2017, FF).

## Declaration of interests

The authors declare no competing interests.

## Figure legends

**Figure S1:**
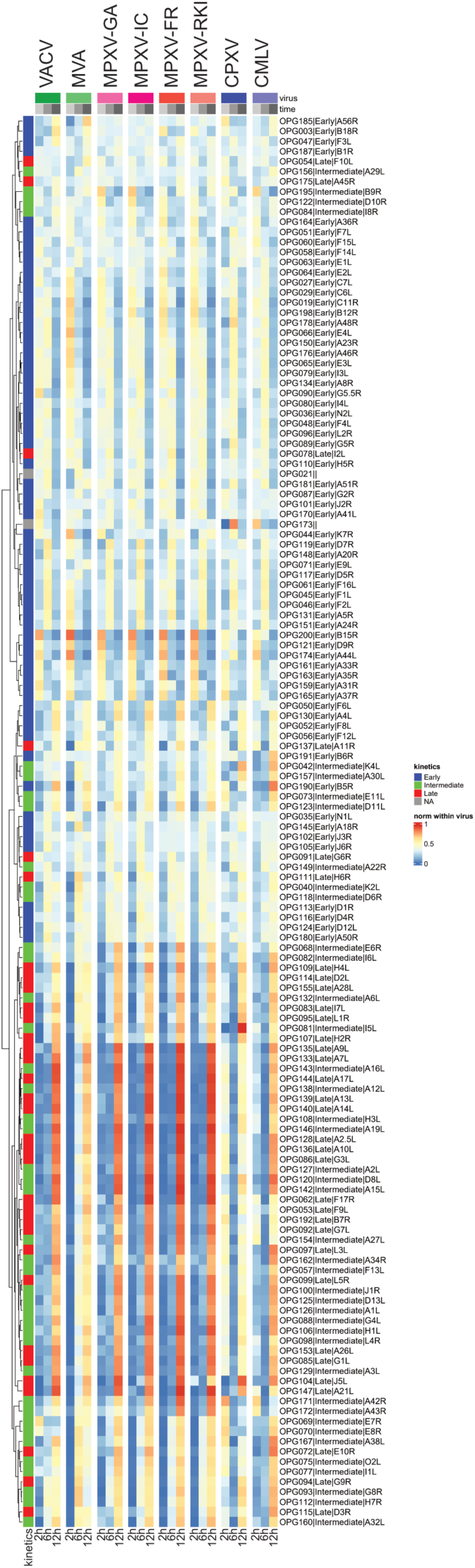
Heatmap of core orthopoxvirus genes. **A)** Heatmap of expression of all core orthopoxvirus genes with early, intermediate and late expression kinetics upon infection with our panel of orthopoxviruses.

**Figure S2:**
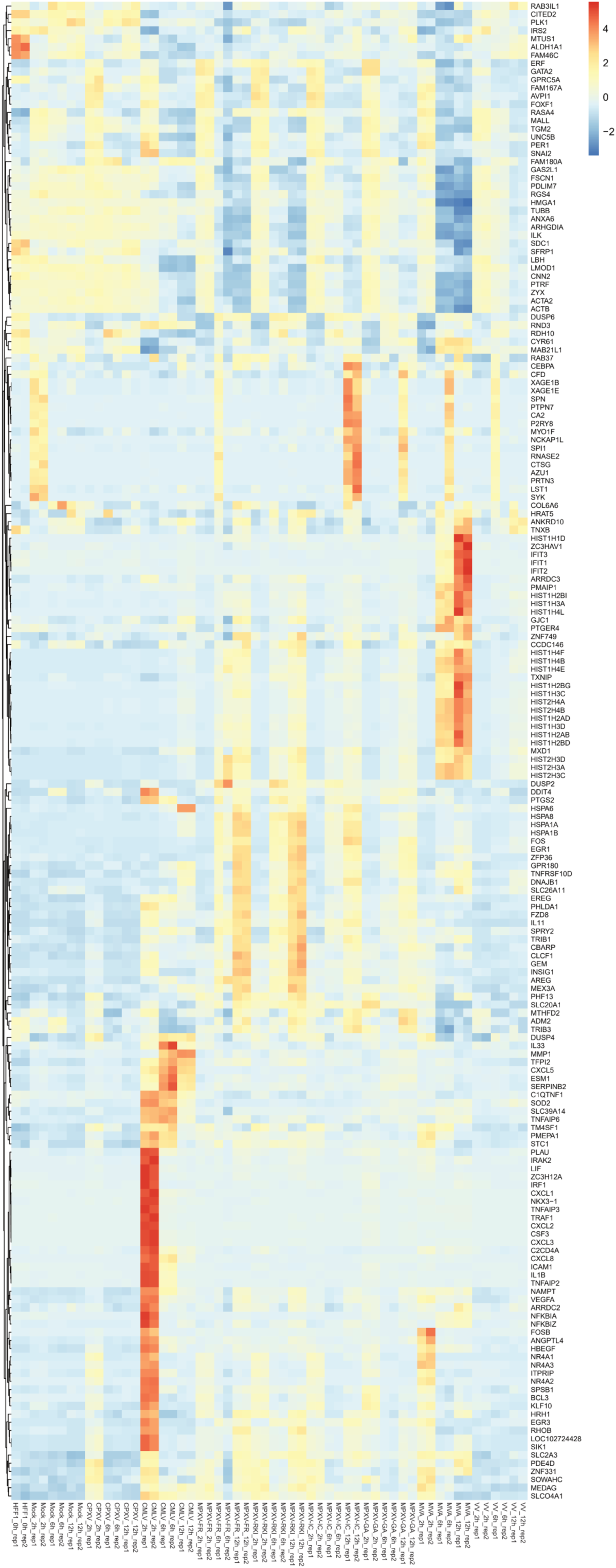
Heatmap of selected host genes. **A)** Heatmap of expression of selected host genes that are differentially regulated by some of the viruses in our panel.

**Figure S3:**
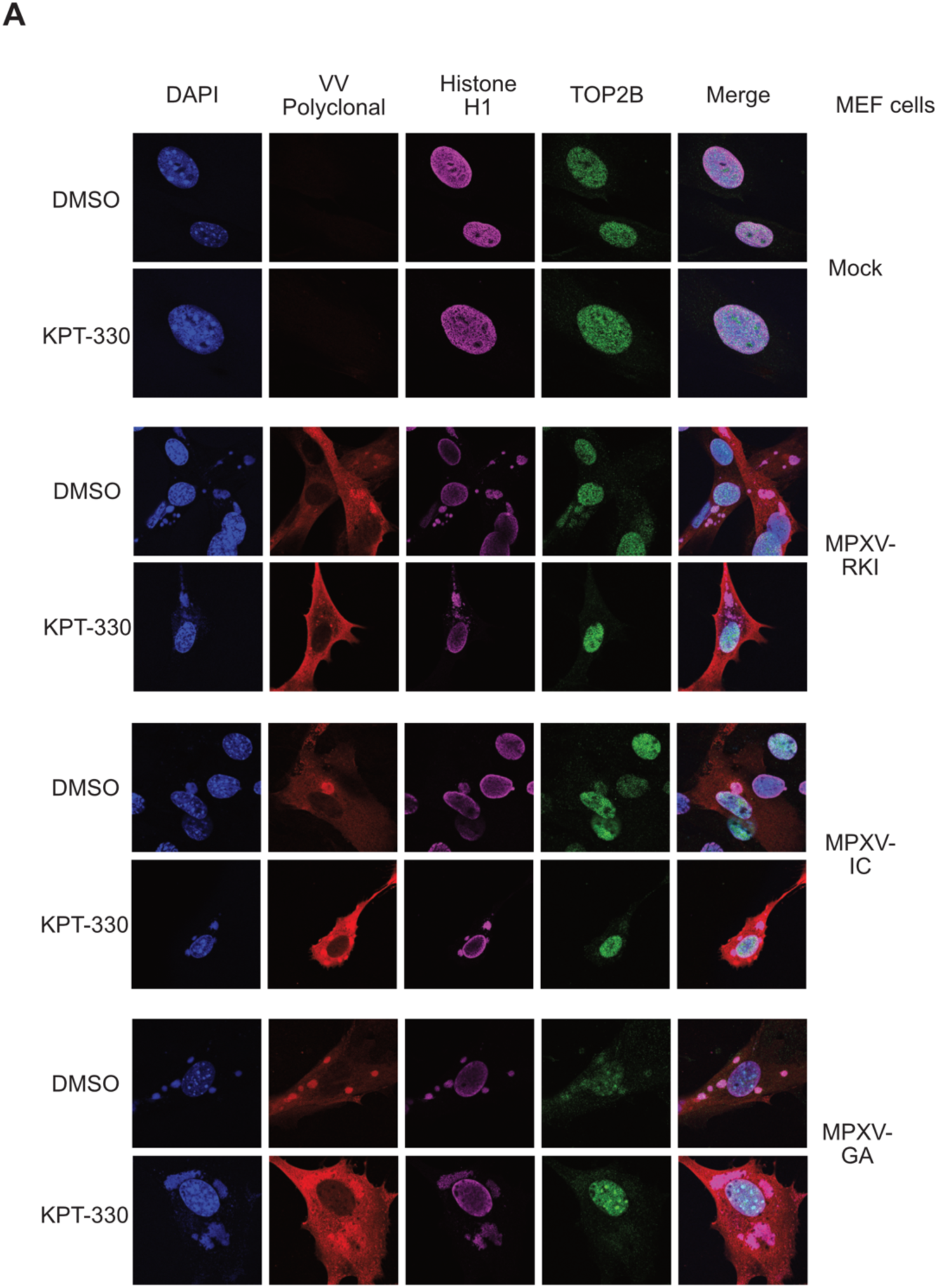
Cytoplasmic transport of Topoisomerase 2 subunits is dependent on the exportin-1 pathway. **A)** Immunofluorescence analysis of MEF cells infected with MPXV strains RKI (MPXV-RKI), Gabun (MPXV-GA) und Ivory Coast (MPXV-IC). Cells were either treated with DMSO (control) or the exportin-1 inhibitor KPT-330 (Selinexor). Cells were fixed and stained with antibodies for Vaccinia Virus (polyclonal), Topoisomerase 2A (TOP2A) and Topoisomerase 2B (TOP2B). Nuclei were counterstained with DAPI.

**Figure S4:**
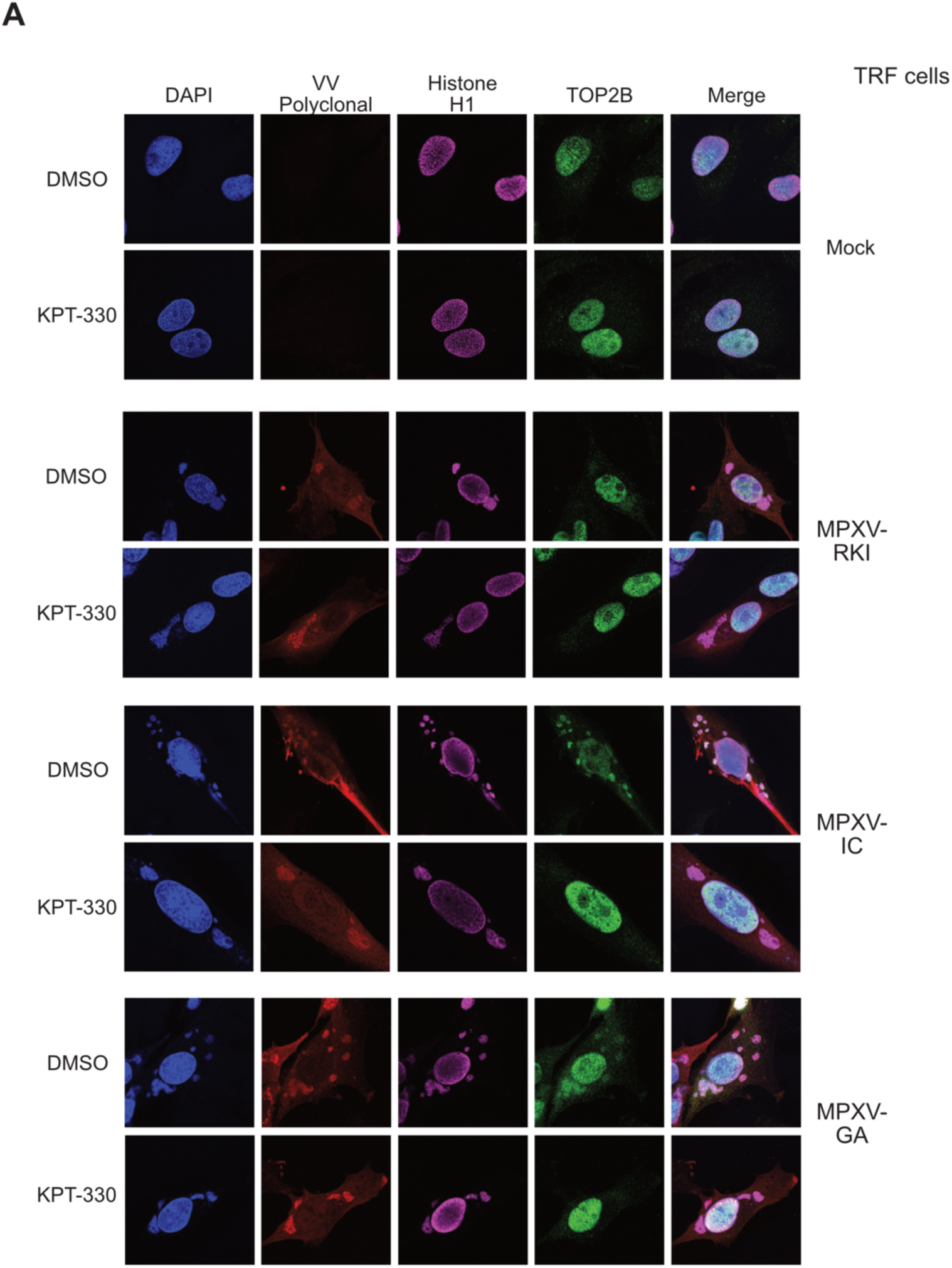
Cytoplasmic transport of Topoisomerase 2 subunits is dependent on the exportin-1 pathway. **A)** Immunofluorescence analysis of Tert-immortalized rhesus fibroblasts (TRF cells) infected with MPXV strains RKI (MPXV-RKI), Ivory Coast (MPXV-IC) and Gabun (MPXV-GA). Cells were either treated with DMSO (control) or the exportin-1 inhibitor KPT-330 (Selinexor). Cells were fixed and stained with antibodies for Vaccinia Virus (polyclonal) histone H1 and Topoisomerase 2B (TOP2B). Nuclei were counterstained with DAPI.

## Notes

### Competing Interest Statement

The authors have declared no competing interest.

